# Historical rice seeds from the 1950s reveal pre-modern genetic structure in *indica* landraces of mainland Southeast Asia

**DOI:** 10.64898/2026.04.19.719523

**Authors:** Koji Numaguchi, Sathya Lim, Chhourn Orn, Yui Higashikubo, Hiroki Saito, Yutaka Sato, Ryo Ishikawa, Rafal Marek Gutaker, Takashige Ishii

**Author notes:** Correspondence: Koji Numaguchi.

## Abstract

Understanding crop population history requires genetic material that predates modern breeding. Here, we analyzed historical seeds of rice landraces assembled in Southeast Asia from 1957 to 1958. Target capture resequencing of coding regions yielded high-quality genomic data from 66 historical accessions (seven from North Vietnam, 30 from South Vietnam, 13 from Cambodia, two from Laos, and 14 from Thailand). When integrated with published rice panels, all historical accessions were assigned to *indica*, and the previously described major regional structure was recovered. Within the historical collection, South Vietnamese, Cambodian, and Thai accessions formed a largely continuous group, whereas North Vietnamese accessions were clearly distinct and comprised two differentiated groups corresponding to the traditional growth seasons (fifth- and tenth-month rice). Notably, the fifth-month rice accessions were assigned to the recently reported Vietnam-I5 cluster. Admixture graph, *f*-statistics, and qpAdm analyses further indicated that Vietnam-I5 is closely related to a China-associated lineage; however, this relationship was not fully explained by sampled *indica, japonica*, or aus proxies. Together, these results show that major components of the present-day *indica* regional structure were already present before modern cultivar replacement, and highlight northern Vietnam as a historical zone of lineage differentiation between Chinese and mainland Southeast Asian rice.

**Highlight:** Genomic analyses of historical *indica* rice seeds reveal that a distinctive northern Vietnamese lineage and a genetic continuum across southern Vietnam, Cambodia, and Thailand predated modern cultivar replacement.

## Introduction

The genetic structure of local crop populations reflects not only biological processes, but also the history of human migration, exchange, and cultural development, and therefore provides a valuable framework for interdisciplinary research linking crop science, anthropology, and archaeology (Fuller *et al*., 2023; Meyer and Purugganan, 2013). However, reconstructing the historical population structure of annual crops such as rice is not straightforward. Most extant plant materials have been reshaped by modern varieties, and the seed bank accessions currently classified as landraces may not fully capture the diversity that existed before large-scale cultivar replacements.

The use of historical DNA offers an opportunity to address this issue. Broadly related to ancient DNA, historical DNA is typically obtained from archived specimens collected decades or a few hundred years ago (Raxworthy and Smith, 2021). Because such material predates modern breeding, it can provide direct access to historical genetic diversity that is no longer represented in extant germplasms. Recent studies have demonstrated the usefulness of historical DNA in reconstructing the past population structures and diversity of cultivated organisms (Alam *et al*., 2025; Burbano and Gutaker, 2023; Forsberg *et al*., 2015; Kistler *et al*., 2025; White *et al*., 2021).

In rice (*Oryza sativa*), large-scale genomic studies have substantially advanced our understanding of domestication and diversification (Alam *et al*., 2025, 2021; Guo *et al*., 2025; Gutaker *et al*., 2020; Jing *et al*., 2023). Recent analyses of extant landraces, particularly *japonica*, have revealed broad geographical structures and dispersal histories (Alam *et al*., 2025, 2021; Gutaker *et al*., 2020). In Vietnam, Higgins *et al*. (2021) further demonstrated a finer structure within local *indica* and *japonica* populations, notably identifying a distinctive outlying *indica* linage, “I5.” These studies provide important insights into the rice population structure in Asia. However, it remains unclear whether these patterns can be detected in historical materials collected before the large-scale replacement of landraces with modern cultivars.

A unique resource for addressing this question is preserved at the Graduate School of Agricultural Science, Kobe University. This collection consists of 1,129 historical rice seeds from Southeast Asian countries, assembled by Professor Hideo Hamada (1899-1994) during the “Southeast Asian Rice Farming Ethnic Culture Research Mission” conducted in 1957–1958 (Matsumoto, 1959). Although seeds lose their germination ability because they are not continuously propagated, DNA can be extracted from seed embryos for genetic analysis (Hiraoka *et al*., 2009). This collection is particularly valuable because it predates both the widespread introduction of modern cultivars after the 1960s and the major political disruptions associated with the Vietnam War and the Cambodian Civil War. Therefore, it provides a rare mid-twentieth-century snapshot of rice diversity in Southeast Asia. The collection included landraces from five regions: North Vietnam, South Vietnam, Cambodia, Laos, and Thailand.

In the present study, we selected 96 accessions spanning the geographical range of this historical collection (hereafter referred to as the Hamada collection) and generated target capture resequencing data for the coding regions. Using previously published landrace datasets as references, we asked whether the broad regional structure inferred from extant *indica* landraces was already present in the 1950s, how historical accessions from mainland Southeast Asia were structured geographically, and whether the distinctive Vietnam-I5 lineage can be traced to pre-modern breeding material. By addressing these questions, we aimed to clarify the historical population structure of *indica* rice in Southeast Asia and place the Hamada collection in the broader context of Asian rice dispersal and lineage formation.

## Materials and Methods

### Materials used for analysis

From the Hamada collection, we selected 96 accessions representing its geographic distribution and usage types, comprising eight accessions from North Vietnam, 33 from South Vietnam, 22 from Cambodia, seven from Laos, and 26 from Thailand (Supplementary Table S1). All seed specimens had lost their germination ability and were stored at room temperature at the Graduate School of Agricultural Science, Kobe University, Japan. For each accession, embryos were excised from 10 seeds, and DNA was extracted using the DNeasy Plant Mini Kit (Qiagen, Hilden, Germany).

Illumina sequencing libraries were prepared using a KAPA HyperPlus Kit (KAPA Biosystems, Wilmington, MA, USA). Each library was barcoded using a single 8-bp NEXTflex adapter (Bioo Scientific, Austin, TX, USA). PCR amplification was performed with PrimeSTAR Max (Takara Bio, Shiga, Japan) under the following conditions: 3 min at 95°C; 8 cycles of 10 s at 95°C, 30 s at 65°C, and 30 s at 72°C; and a final extension for 5 min at 72°C. The resulting libraries were used for the target capture experiment.

For target capture, RNA probes were designed using MyBaits Custom Design Kit 1–20K probes (Arbor Biosciences, Ann Arbor, MI, USA). In total, 16,414 loci were targeted, including 16,402 coding and 12 simple sequence repeat (SSR) loci, and 16,462 RNA probes were designed (16,402 for coding and 60 for SSR loci). The probe sequences are publicly available from Zenodo (https://doi.org/10.5281/zenodo.19434769; accessed on April 6, 2026). Genomic DNA libraries were pooled in equimolar amounts and subjected to hybridization capture according to the manufacturer’s instructions. Captured libraries were amplified with PrimeSTAR Max under the following conditions: 3 min at 95°C; 14 cycles of 10 s at 95°C, 30 s at 65°C, and 30 s at 72°C; and a final extension for 5 min at 72°C. Sequencing was performed on an Illumina HiSeq 4000 platform with paired-end 100-bp reads. The accession numbers for all sequence data generated in this study are listed in Supplementary Table S1.

For integrative analyses, we downloaded the following whole-genome sequencing (WGS) datasets from public databases: 69 accessions from the World Rice Core Collection (WRC) and 50 accessions from The Rice Core Collection of Japanese Landraces (JRC) (Tanaka *et al*., 2020a; 2020b); 337 accessions belonging to six *indica* subpopulations defined by PAM-silhouette clustering (Kd = 6) in Gutaker *et al*. (2020) [CHNO (China/Taiwan), KHM (Cambodia), IDN3 (Indonesia), IND2 (India), LAO (Laos), and THA (Thailand/Malaysia)]; and 64 accessions assigned to the I1–I4 subpopulations (depth > 4) and 38 accessions assigned to the I5 subpopulation from Higgins *et al*. (2021). Accession numbers for the WGS datasets are listed in Supplementary Table S2.

### Read processing and SNP calling

#### Hamada collection

Target capture reads from the Hamada collection (Supplementary Table S1) were processed following workflows commonly used for historical DNA data (Latorre *et al*., 2020). Adapter trimming and merging of overlapping paired-end reads were performed using AdapterRemoval v.2.3.1 (Schubert *et al*., 2016). This step was included to improve the per-base quality score, adapter detection and overall read utilization of historical DNA, which often contains very short fragments. Quality trimming was performed using Trimmomatic v.0.39 (Bolger *et al*., 2014) with the following parameters: LEADING:20, TRAILING:20, SLIDINGWINDOW:4:20, and MINLEN:35. The merged and unmerged paired-end reads were combined for downstream analyses.

The processed reads were mapped to the *indica* Guangluai-4 reference genome (Guo *et al*., 2025) using BWA aln (Li and Durbin, 2009) with the disabled seed (-l 1024) option. SAM/BAM file processing was performed using Samtools v.1.9 (Li *et al*., 2009). Duplicate reads were removed using DeDup v.0.12.8 (Peltzer *et al*., 2016). The DNA damage patterns were assessed using mapDamage v.2.3.0 (Jónsson *et al*., 2013). Although this tool can be used to downscale the quality scores of putatively damaged bases, we did not apply it because our samples showed little evidence of DNA damage.

SNP calling was performed using GATK v.4.5.0.0 (Van der Auwera and O’Connor, 2020) with the HaplotypeCaller, GenotypeGVCFs, and SelectVariants functions. Called SNPs were filtered using VariantFiltration with the following criteria: QualByDepth (QD) < 2.0, FisherStrand (FS) > 60.0, StrandOddsRatio (SOR) > 3.0, MappingQuality (MQ) < 30.0, MappingQualityRankSumTest (MQRunkSum) < −12.5, and ReadPosRankSumTest (ReadPosRankSum) < −8.0 Details for these parameters are provided in the GATK documentation (https://gatk.broadinstitute.org/hc/en-us/articles/360035890471-Hard-filtering-germline-short-variants; accessed on July 1, 2026). Using VCFtools (Danecek *et al*., 2011), genotypes with minDP < 4 were treated as missing, and samples with a missing genotype rate > 0.5 were removed. This dataset is hereafter referred to as hamada-VCF.

#### Public WGS data

For publicly available WGS data (Supplementary Table S2), reads were quality-trimmed using Trimmomatic as described above and mapped to the Guangluai-4 reference genome using the BWA mem (Li, 2013).

To generate an SNP set comparable to the Hamada target capture data, we first filtered the hamada-VCF using VCFtools to retain SNPs with a missing genotype rate < 0.5. We then used pysam (https://github.com/pysam-developers/pysam; accessed on March 24, 2026) to extract paired-end reads overlapping these SNP positions from public WGS data (BAM files). Duplicate reads were removed using GATK MarkDuplicates and SNP calling was performed as described above. VariantFiltration was applied using the following criteria: QD < 2.0, FS > 60.0, SOR > 3.0, MQ < 40.0, MQRankSum < −12.5, and ReadPosRankSum < −8.0.

#### Panel construction

Three analytical panels were prepared for this study (Panels 1–3; Table 1).

**Table 1.** Summary of analytical panels used in this study.

| Panel | Description<sup>1</sup> | Total number of samples |
| --- | --- | --- |
| Panel 1 | All accessions, including the *japonica* (60), *indica* (544), and aus (20) groups | 624 |
| Panel 2 | Hamada collection accessions only | 66 |
| Panel 3 | Indica-focused panel including the Hamada collection (66) and published *indica* accessions (478) | 544 |

<sup>1</sup> Numbers in parentheses indicate the number of samples in each group.

Panel 1 shows the 624 accessions used in the present study. Hamada-VCF and publicly available VCF datasets were merged, and variants were filtered using VCFtools with --remove-filtered-all and --max-missing 0.8. Imputation was performed using Beagle 5.5 (Browning *et al*., 2021) with the default settings. Subsequently, SNPs with a minor allele frequency < 0.05 were removed using PLINK (Purcell *et al*., 2007), and linkage disequilibrium (LD)-based pruning was performed with --indep-pairwise 10kb 1 0.8.

Panel 2 comprises the 66 Hamada collection accessions that were retained after quality filtering. Hamada-VCF was filtered using VCFtools with --remove-filtered-all and --max-missing 0.8, followed by imputation with Beagle 5.5. SNPs with a minor allele count < 2 were removed using PLINK and LD pruning was performed with --indep-pairwise 10kb 1 0.8. This panel was used for the analysis of the Hamada collection alone and for SNP annotation using SnpEff (Cingolani *et al*., 2012b).

Panel 3 comprises 544 accessions obtained by merging the Hamada collection with *indica* accessions from public datasets. After merging, the filtering, imputation, and LD-pruning steps used in Panel 1 were applied.

#### Population structure analysis

Principal component analysis (PCA) was performed using PLINK on the LD-pruned SNP sets in each panel. PAM-silhouette-based clustering was conducted following Gutaker *et al*. (2020), using pairwise genetic distance matrices (1-IBS). NeighborNet analysis was conducted using IBS distance-based input data. NEXUS files were generated from the VCF data of panels 1–3 using gdsfmt, SNPRelate, SeqArray, and Phangorn (Schliep, 2011; Zheng *et al*., 2017, 2012). NeighborNet analysis and visualization were performed using the SplitsTree App (Huson and Bryant, 2006). ADMIXTURE analysis was performed using ADMIXTURE (Alexander *et al*., 2009). Multiple *K* values were examined for each panel and the principal results are presented. Although the cross-validation error was the lowest at *K* = 3 (CVE = 0.645), *K* = 6 (CVE = 0.739) was also considered because it provided a more interpretable population structure and was that aligned more closely with the PCA and NeighborNet results. Pairwise genetic differentiation was estimated using the Weir _–_ Cockerham *F*_ST_. Calculations were performed using the VCFtools in Panel 2 with groups defined by country or region.

#### Computation of *f*-statistics

To evaluate the genetic relationships between the Hamada collection and other subpopulations, outgroup *f*_3_ and *f*_4_ statistics were calculated. Input files were prepared by adding 19 accessions of *O. barthii* (Alam *et al*., 2025; Choi *et al*., 2019; Gutaker *et al*., 2020; Stein *et al*., 2018) (Supplementary Table S2) as the outgroup to Panels 1 and 3, and by specifying assignment to Kd = 9 clusters (*japonica*, aus, Vietnam-I5, China, India, Indonesia, Cambodia, Thailand, and Laos) for Panel 1 and Kd = 7 clusters (Vietnam-I5, China, India, Indonesia, Cambodia, Thailand, and Laos) for Panel 3 in PLINK format (bed/bim/fam). Calculations were performed using ADMIXTOOLS 2.0.4 (Maier *et al*., 2023). For the outgroup *f*_3_, *f*_3_(*O. barthii*; X, Y) was calculated. For *f*_4_, we calculated *f*_4_(*O. barthii*, X; Y, Z). Analyses plotting f-statistics against PC1, as well as analyses those ordering Vietnam-I5 accessions according to PC2, were based on these input files. Correlations between the *f*-statistics and the PCs were evaluated using Pearson’s correlation coefficient.

### Admixture graph construction and TreeMix analysis

To infer the formation history of *indica* subpopulations defined by PAM-silhouette clustering (Kd = 7) (Panel 3), admixture graphs were constructed using the qpGraph function in Admixtools 2.0.4. PLINK-formatted input files were prepared from Panel 3 with assignment to the seven clusters (Vietnam-I5, China, India, Indonesia, Cambodia, Thailand, and Laos). The topology searches were performed using the find_graphs function with stop_gen = 100 and stop_gen2 = 20. Candidate graphs were sorted by score, and the maximum residual |Zmax∣ was calculated for the top 10 candidates. For each numadmix value, the graph with the smallest |Zmax| value was selected. We increased numadmix sequentially from 0 and selected the first graph satisfying ∣Zmax∣ < 3 as the best-fitting graph.

For TreeMix analysis, population-level allele frequencies generated by PLINK (--freq, producing frq.strat) were converted into TreeMix format using the developer-provided script plink2treemix.py. Analyses were conducted using migration edges (m) =0 to m=2. The block-size parameter -k, which specifies the number of SNPs grouped into each block, was set to 500.

#### qpAdm analysis

To test whether Vietnam-I5 could be represented by sampled proxies for cultivated rice sources, we used qpAdm as implemented in ADMIXTOOLS 2.0.4. Among accessions assigned to Vietnam-I5, WRC22, WRC24, and WRC44 were excluded from qpAdm analysis because they exhibited exceptionally high *japonica* affinity. The remaining Vietnam-I5 accessions were divided into Low-PC2 and High-PC2 groups based on PCA and *f*-statistic patterns, and each group was treated as a target population. For each target, we tested four source-proxy models: (1) China only, (2) *japonica* only, (3) aus only, (4) *japonica* + aus, (5) China + *japonica*, (6) China + aus, and (7) China + *japonica* + aus. In these models, the listed groups were included as separate source proxies, and allele frequencies were not pooled across groups. Here, the target population and candidate source proxies constituted the “left” population set. Populations from Cambodia, Indonesia, India and Thailand were used as “right” reference populations to distinguish the ancestry relationships of the target and source proxies. These groups were selected as geographically and genetically differentiated cultivated *indica* reference populations, and were not included as target populations or source proxies. For sensitivity analysis, we repeated the qpAdm tests with *O. barthii* added to the right population set as the outgroup.

The qpAdm tests whether the *f*_4_ statistic pattern of the target relative to the right populations can be represented as a linear combination of the corresponding *f*_4_ patterns of the source proxies. The null hypothesis of a qpAdm model is that the target population is a mixture of the specified source proxies, in the sense that the source proxies fully account for the shared drift between the target and right populations. For a model with k source proxies, this corresponds to testing whether the target can be explained by a rank k - 1 model. Here, the rank summarizes the number of independent ancestry relationships required to explain the *f*_4_ statistic patterns between the left and right populations. Model *P*-values indicate the fit of the specified source model, and *P* < 0.05 was interpreted as rejection of the null hypothesis. The admixture coefficients were interpreted only for models that were not rejected and yielded coefficients within the feasible range of 0–1.

## Results

### Sequencing read statistics for the Hamada collection

A total of 96 accessions of the Hamada collection (eight from North Vietnam, 33 from South Vietnam, 22 from Cambodia, seven from Laos, and 26 from Thailand) were sequenced using target capture of 16,402 coding and 60 SSR loci. The number of reads obtained per accession ranged from 185,535 (Cam-6) to 18,556,029 (Tha-484) (Supplementary Table S1). Read mapping rates were consistently high across all accessions, ranging from 90.1% (ViS-937) to 94.7% (Lao-187). Duplicate read rates varied widely, from 8.8% (Tha-473) to 83.0% (Lao-185) (Supplementary Table S1).

To evaluate whether the reads exhibited damage patterns characteristic of historical DNA, we analyzed all 96 samples using mapDamage v.2.3.0 (Jónsson *et al*., 2013). Because broadly similar patterns were observed across all accessions, we present the results for ViS-956 as a representative example (Supplementary Fig. S1). There was little evidence of substantial C-U deamination. The read length distribution displayed a pronounced peak at 100 bp, corresponding to unmerged paired-end (PE) 100 reads, together with a continuous distribution from 35 bp (the lower filtering threshold in Trimmomatic) to approximately 190 bp, which was consistent with the merged paired-end reads generated by AdapterRemoval 2 (Supplementary Fig. S1).

After excluding samples with a missing genotype rate > 0.5, we retained 66 high-quality samples (seven accessions from North Vietnam, 30 from South Vietnam, 13 from Cambodia, two from Laos, and 14 accessions from Thailand indicated as “Pass” in Supplementary Table S1). To characterize the genomic distribution of the single nucleotide polymorphisms (SNPs) used in downstream analyses, we annotated the quality-filtered SNP set from these 66 samples, using SnpEff (Cingolani *et al*., 2012a), with genotypes called at a minimum depth of ≧ 4. Most SNPs (83.6%) were located in introns (Supplementary Table S3). This pattern indicated that targeted resequencing successfully captured not only the intended exonic targets, but also adjacent intronic regions, thereby yielding abundant polymorphic sites (n > 10,000) for population structure analyses.

### The Hamada collection consists exclusively of *indica* accessions

To determine whether the Hamada collection accessions belonged to the *japonica, indica*, or aus groups, we constructed a merged panel of 624 accessions comprising the Hamada collection and public *japonica, indica*, and aus accessions (Panel 1; Table 1). For the Hamada collection, the 66 accessions listed above were retained for downstream analyses to ensure data quality (“Pass” in Supplementary Table S1). Using 13,362 SNPs that passed the filtering, principal component analysis (PCA) clearly separated the major rice groups; *japonica*, aus and *indica* (Fig. 1A).

**Fig. 1.**
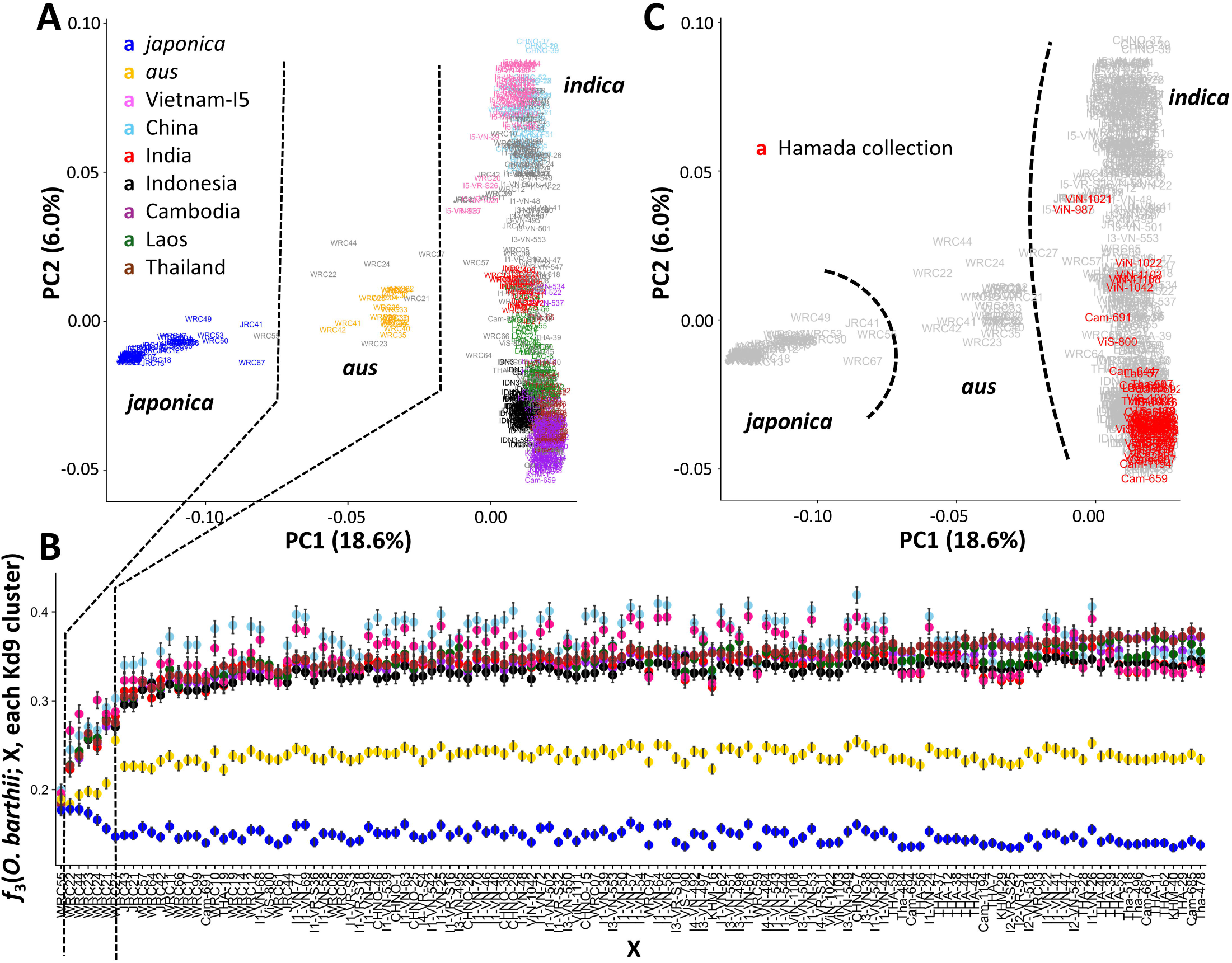
Overall population structure of Panel 1 and cultivar-group affiliation of the Hamada collection accessions. **A** Principal component analysis (PCA) of all 624 accessions used in this study, including *japonica*, aus, and *indica* groups. Colors indicate discrete subpopulations defined by PAM-silhouette based clustering at Kd = 9. **B** Outgroup *f*_3_ analysis of accessions not assigned to the subpopulations shown in **A**. Bars indicate standard errors estimated using block jackknife. Dotted lines indicate tentative boundaries associated with *japonica-aus* and *aus-indica* transitions. **C** PCA plot highlighting the Hamada collection accessions in Panel 1. All the Hamada collection accessions fell within the *indica-* related cluster.

To further characterize the population structure of Panel 1, we applied partitioning around medoids (PAM)-silhouette clustering based on a pairwise genetic distance matrix, following Gutaker *et al*. (2020). Across Kd = 2–10, the major groupings were consistently recovered, and at Kd = 9 the clustering broadly reproduced the previously reported *indica* structure of Gutaker *et al*. (2020), while also recovering the Vietnam-I5 linage defined by Higgins *et al*. (2021) (Fig. 1A; Supplemenrary Fig. S2).

For accessions left unassigned by Kd = 9 clustering, we calculated the outgroup *f*_3_ statistics, *f*_3_(*Oryza barthii*; each unassigned line, each Kd9 cluster) and ordered the unassigned lines using principal component (PC) 1 (Fig. 1B) to measure the strength of their genetic affinity to discrete genetic clusters. This approach was used complementarily to define the *japonica*-aus and aus-*indica* boundaries and to position accessions that were not clearly assigned to discrete PAM-silhouette clusters. Several accessions with lower PC1 values, namely WRC55, WRC22, WRC44, WRC23, WRC24, WRC21, and WRC27, showed elevated drift sharing with *japonica* (WRC55) and aus (the others). In contrast, unassigned accessions with higher PC1 values exhibited greater drift sharing with *indica* clusters. Based on this pattern, we treated the latter as belonging to the *indica* lineage.

All 66 retained Hamada accessions were assigned to *indica*-associated clusters (Panel 1), indicating that the Hamada collection consisted exclusively of *indica* accessions (Fig. 1C).

### Historical population structure of mainland Southeast Asian *indica* landraces in the Hamada collection

We constructed a panel restricted to the Hamada collection (Panel 2; Table1) to examine the population structure of mainland Southeast Asian *indica* landraces before the widespread replacement of traditional cultivars with modern elite varieties. The PCA based on 10,136 SNPs from 66 accessions, revealed that most accessions formed a single major cluster (Fig. 2A). However, two North Vietnamese accessions (ViN-987 and ViN-1021; classified as “fifth-month rice,” Supplementary Table S1), one Cambodian accession (Cam-691), and one South Vietnamese accession (ViS-800) were displaced primarily along PC1, whereas five North Vietnamese accessions (ViN-1108, ViN-1103, ViN-1022, ViN-1042, and ViN-1111; classified as “tenth-month rice,” Supplementary Table S1) were displaced primarily along PC2 (Fig. 2A). The fifth and tenth month rice cultivation correspond to two traditional rice-growing seasons in northern Vietnam (Hamada, 1965).

**Fig. 2.**
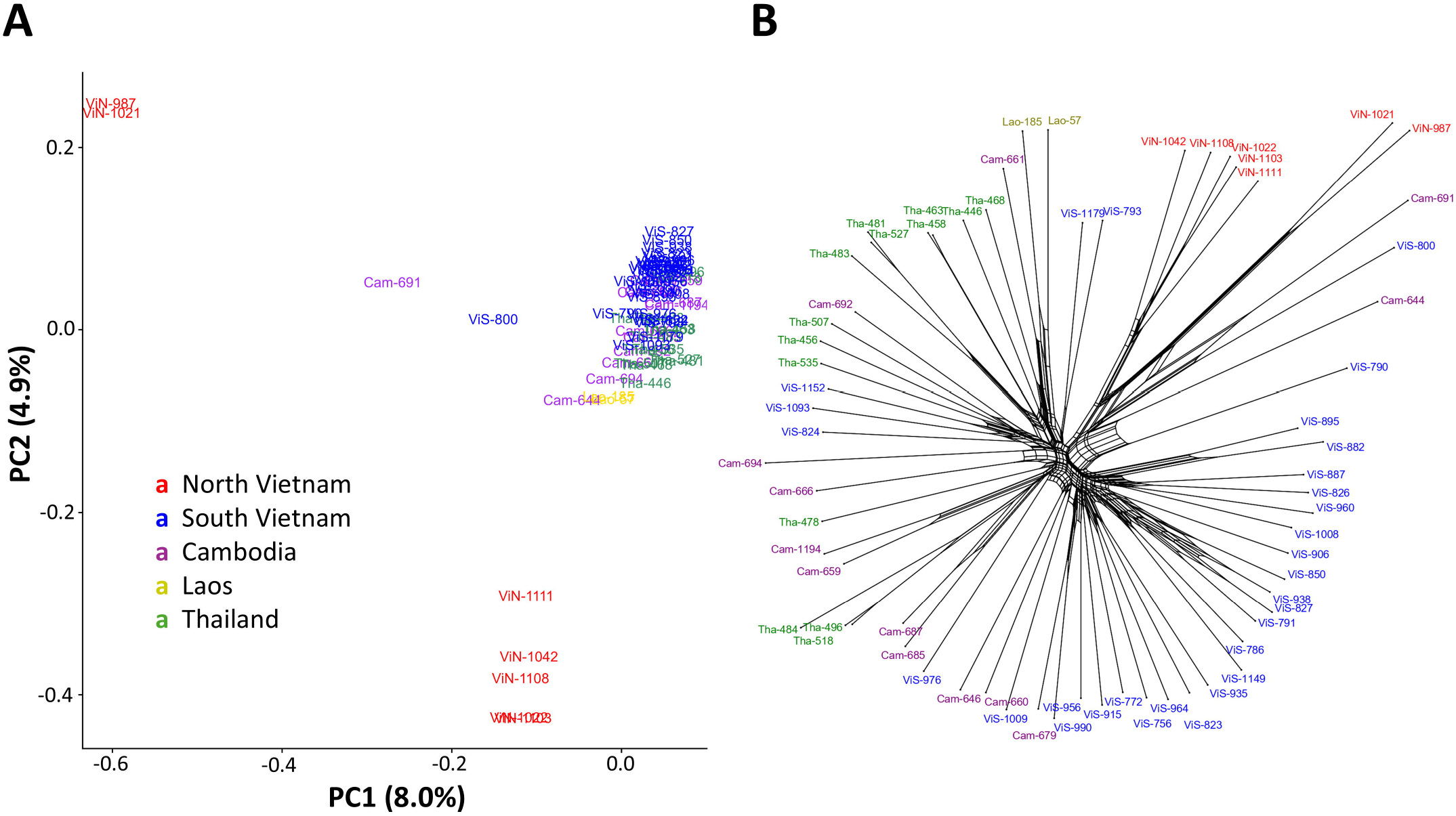
Pre-modern population structure inferred from the Hamada collection accessions (Panel 2). **A** Principal component and **B** NeighborNet analyses for Panel 2 consist of 66 Hamada collection accessions. Colors were assigned to each group according to their geographic origin.

The NeighborNet analysis broadly recapitulated this pattern. In addition to the nine outlying accessions identified by the PCA, one Cambodian accession (Cam-644) and one South Vietnamese accession (ViS-790) occupied displaced positions within the network (Fig. 2B). Most of the remaining accessions, including those from South Vietnam (ViS), Cambodia (Cam), Thailand (Tha), and Laos (Lao), formed a broad central assemblage with some level of geographic structuring, but were connected by a complex reticulate network, suggesting substantial genetic exchange among regional landraces. The overall star-like topology further indicated a limited hierarchical structure among historical accessions (Fig. 2B).

ADMIXTURE analyses supported both the distinctiveness of the North Vietnamese accessions and the genetic proximity profiles observed in many South Vietnamese/Cambodian/Thai accessions (Supplementary Fig. S3). The relationship graph obtained using the TreeMix approach supports these patterns, but the residual fit between the North and South Vietnamese groups was improved slightly when a migration edge between them was added (Supplementary Fig. S4).

These patterns were also supported by the pairwise *F*_ST_ estimates (Table 2). Cambodian rice exhibited very low differentiation from South Vietnamese (*F*_ST_ = 0.042) and Thai (*F*_ST_ = 0.040) rice, whereas North Vietnamese rice was more strongly differentiated from Laos (*F*_ST_ = 0.290), South Vietnamese (*F*_ST_ = 0.230), and Thai (*F*_ST_ = 0.238) rice. Together, these results indicate a distinctive North Vietnamese group, alongside a broader Cambodia-South Vietnam-Thailand continuum of historically connected *indica* landraces.

**Table 2.** Weir and Cockerham *F*_ST_ estimates among geographic groups of the Hamada collection.

|  | Cambodia | Laos | North Vietnam | South Vietnam | Thailand |
| --- | --- | --- | --- | --- | --- |
| Cambodia |  |  |  |  |  |
| Laos | 0.132 |  |  |  |  |
| North Vietnam | 0.178 | 0.290 |  |  |  |
| South Vietnam | 0.042 | 0.236 | 0.230 |  |  |
| Thailand | 0.040 | 0.193 | 0.238 | 0.101 |  |

### *Indica* population structure reproduces geographic subdivision and the distinctiveness of North Vietnamese accessions

In the previous section, an analysis of the Hamada collection revealed the historical population structure of mainland Southeast Asian *indica* landraces. To place these historical accessions in a broader geographic and temporal context, we constructed an *indica*-wide panel (Panel 3; Table 1) by merging our sequence data with previously published datasets (Gutaker *et al*., 2020; Higgins *et al*., 2021; Tanaka *et al*., 2020a; 2020b) and performed population structure analyses.

Using 12,227 SNPs, PCA and PAM-silhouette clustering showed that, at Kd = 7, the major *indica* structure broadly reproduced the subdivision reported by Gutaker *et al*. (2020), while also recovering the Vietnam-I5 lineage defined by Higgins *et al*. (2021) (Fig. 3A; Supplementary Figs S5, S6). Most Hamada accessions were assigned to the Cambodian-, Thai-, or Laos-related clusters. South Vietnamese, Cambodian, and Thai accessions from the Hamada collection did not form fully discrete groups, but instead overlapped extensively, consistent with the mixed regional structure already observed in the Hamada-only analyses (Fig. 3A; Supplementary Fig. S6). Among the North Vietnamese accessions, the two fifth-month rice accessions (ViN-987 and ViN-1021) were assigned to the Vietnam-I5 cluster, whereas the five tenth-month rice accessions (ViN-1108, ViN-1103, ViN-1022, ViN-1042, and ViN-1111) were not assigned to any Kd = 7 cluster and were positioned near the Indian cluster in PCA space (Fig. 3B; Supplementary Fig. S6). The four outlier accessions identified in the Hamada-only analysis (Cam-644, Cam-691, ViS-790, and ViS-800) were positioned between the Indian, mainland Southeast Asian, and Indonesian clusters (Supplementary Fig. S6).

**Fig. 3.**
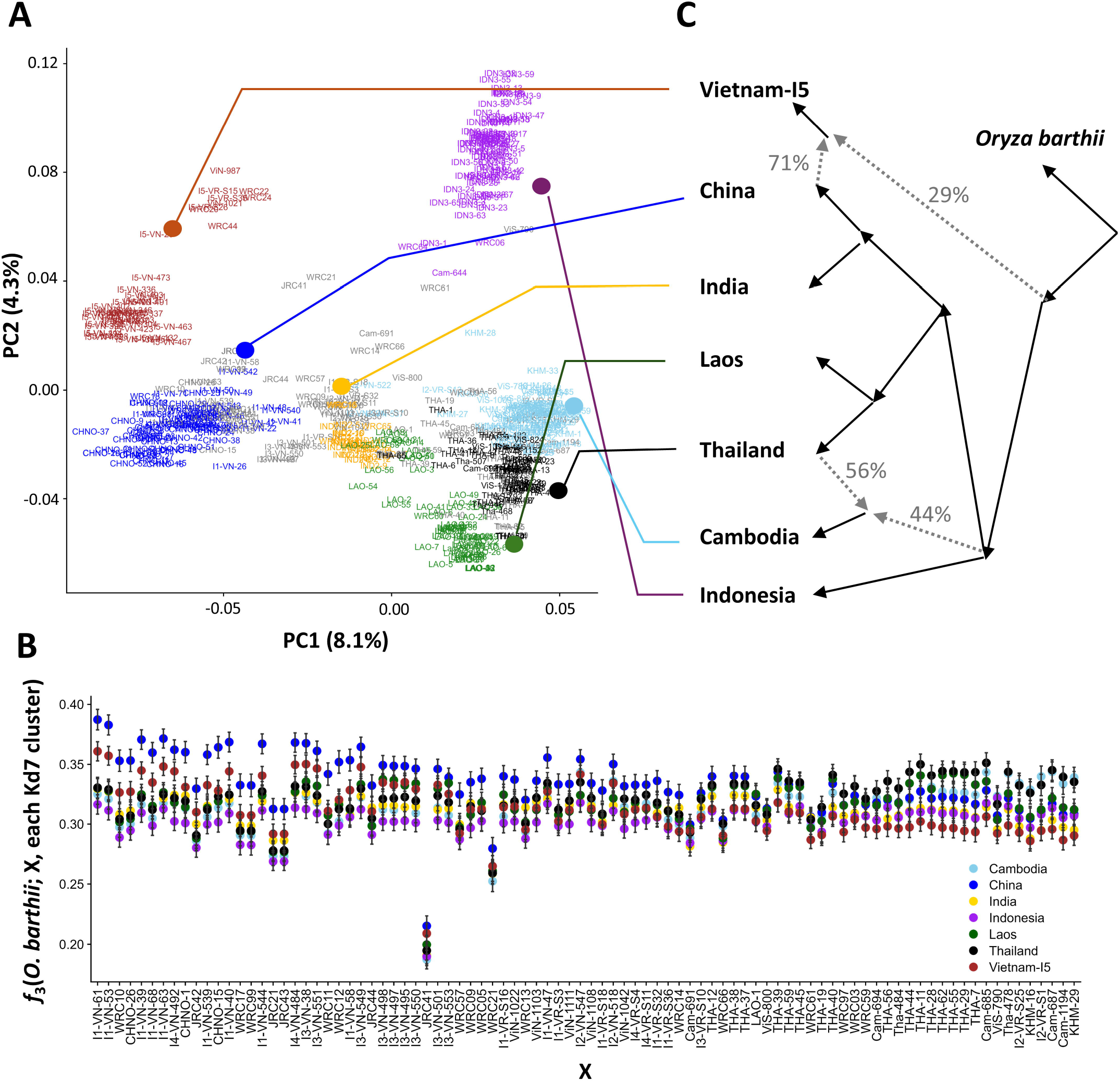
*indica-wide* population structure of Panel 3 and admixture relationships among inferred subpopulations. **A** Principal component analysis of the 544 *indica* accessions shown in Panel 3. Colors indicate discrete subpopulations defined by PAM-silhouette-based clustering at Kd = 7. **B** Outgroup *f*_*3*_ analysis of accessions not assigned to the subpopulations shown in **A**. Bars indicate standard errors estimated using block jackknife. **C** Best-fitting admixture graph inferred from Kd = 7 subpopulations. The dotted arrows indicate the inferred admixture proportions between the ancestral components.

To further characterize the accessions that were not assigned to discrete clusters (Kd = 7), we calculated the outgroup *f*_3_ statistics, *f*_3_ (*O. barthii*; each unassigned accession, each Kd7 cluster), and ordered the accessions by PC1 (Fig. 3B). Broadly, drift sharing with the Chinese and Vietnam-I5 clusters tended to decrease along PC1, whereas drift sharing with the Cambodian, Laos, and Thai clusters tended to increase. By contrast, no clear monotonic trend was observed for the Indian and Indonesian clusters.

We then performed admixture graph analyses for the Kd = 7 clusters using qpGraph in ADMIXTOOLS 2 (Maier *et al*., 2023). Running the find_graphs function with the number of admixture events fixed from 0 to 2 identified a graph with two admixture events and a worst |Z| = |−2.82| < 3 (Fig. 3C). This result was broadly consistent with TreeMix (Pickrell and Pritchard, 2012) analyses (Supplementary Fig. S7). In particular, the best-fitting graph indicated that Vietnam-I5 could be modeled as a China-related lineage with an additional unknown component. This interpretation is consistent with the outgroup *f*_3_ results, which indicate that unassigned accessions with lower PC1 values share more drift with the Chinese and Vietnam-I5 clusters than those with higher PC1 values. In addition, the Cambodian cluster was best modeled as carrying ancestry related to both Thai and Indonesian clusters, in agreement with the broad pattern reported by Gutaker *et al*. (2020).

### Vietnam-I5 carries China-related ancestry but cannot be fully explained by ancestries of cultivated groups sampled in this study

To examine the ancestry components distinguishing the Chinese and Vietnam-I5 clusters from the other *indica* groups, we performed a series of *f*-statistics analyses (Fig. 4). The outgroup *f*_3_ statistics, *f*_3_(*O. barthii*; each accession assigned to the Kd7, *japonica*/aus cluster), were plotted against PC1. Shared drift with both *japonica* and aus decreased broadly monotonically along PC1 and showed significant negative correlations with PC1 (R = −0.90 and −0.63, respectively; *P* < 2.2 × 10^−16^; Fig. 4A, B), with the stronger relationship observed for *japonica*. We also calculated Z[*f*_4_(*O. barthii*, each accession assigned to Kd7; *japonica* cluster, aus cluster)], which exhibited a significant positive correlation with PC1 (R = 0.43, *P* < 2.2 × 10^−16^; Fig. 4C). Together, these results suggest that the separation of the Chinese and Vietnam-I5 clusters from the remaining *indica* clusters along PC1 is associated with relatively greater shared drift with *japonica* and aus. Notably, the Indian cluster showed particularly high *f*_4_ Z scores, consistent with a stronger aus-related affinity than other *indica* groups (Fig. 4C).

**Fig. 4.**
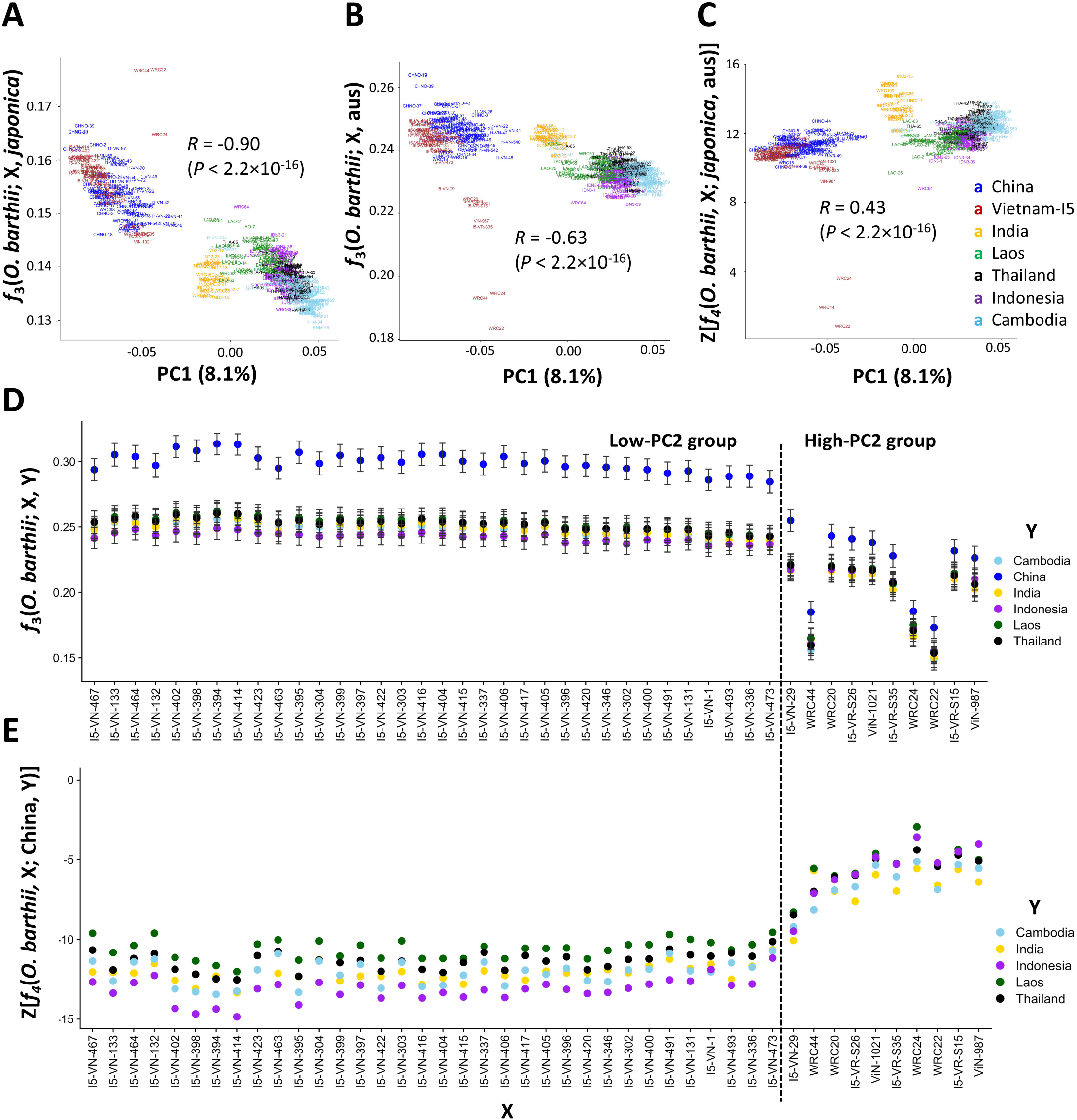
Vietnam-IS and China-related accessions show affinities with *non-indica* ancestry components. **A, B** Outgroup *f*_*3*_ analyses of accessions assigned to Kd =7 clusters {Panel 3) with *japonica* and aus clusters, respectively, plotted against PCl. CZ scores of *f*_*4*_ statistics indicating the relative affinity of each accession assigned to Kd = 7 clusters for *japonica* and aus. **D** Outgroup *f*_*3*_ analysis of accessions belonging to the Vietnam-IS cluster with each Kd = 7 cluster, ordered by PC2. Bars indicate standard errors estimated using block jackknife. **E** Z scores of *f*_4_ statistics indicating the relative affinity of each Vietnam-IS accession to the China cluster and the other Kd = 7 clusters, ordered by PC2. The dotted line indicates the tentative boundary between Low-PC2 and High-PC2 groups.

The Vietnam-I5 cluster was further distinguished from the Chinese cluster primarily along PC2 and formed a relatively broad cline along this axis (PC2 = −0.07 to 0.11; Fig. 3A). To assess how affinity to the other *indica* clusters varies within Vietnam-I5, we calculated outgroup *f*_3_ statistics, *f*_3_(*O. barthii*; each Vietnam-I5 accession, each Kd7 cluster), and ordered the accessions by PC2. Vietnam-I5 accessions consistently shared the most drift with the Chinese cluster, but this value declined sharply around the transition marked by I5-VN-29 (Fig. 4D). We then calculated Z[*f*_4_(*O. barthii*, each Vietnam-I5 accession; Chinese cluster, each other *indica* cluster)], again ordered by PC2. In parallel with the *f*_3_ pattern, the Z-scores increased markedly toward the high-PC2 end across all five tested *indica* clusters (Fig. 4E).

Based on the abrupt shift in *f*-statistics around I5-VN-29, we defined accessions located on the high-PC2 side of this transition as the High-PC2 group and those on the low-PC2 side as the Low-PC2 group (dotted line in Fig. 4D, E). Three Vietnam-I5 accessions (WRC22, WRC24, and WRC44) were excluded from these analyses because they showed extremely high *japonica* affinity, consistent with the results shown in Fig. 1B. We then used qpAdm to test whether the Low-PC2 and High-PC2 subsets of Vietnam-I5 could be adequately modeled using one-, two-, or three-source models involving the Chinese, *japonica*, and aus clusters. For the Low-PC2 subset, the one-source models (China only, *japonica* only, and aus only), the two-source models (*japonica* + aus, China + *japonica*, and China + aus), and the three-source model (China + *japonica* + aus) were all rejected (Table 3). For the High-PC2 subset, the *japonica* only model was not rejected when *indica* populations alone were used as right-set references, whereas the other models were either rejected or yielded infeasible admixture coefficients (Table 3). However, when *O. barthii* was added to the right population set, all models for both the Low- and High-PC2 subsets were strongly rejected (Supplementary Table S4). The contrasting results across right-set configurations indicate that the non-rejection of *japonica*-only model for the High-PC2 subset was not robust across analyses, although this subset appeared to show stronger *japonica*-related affinity than the Low-PC2 subset (Table3; Supplementary Table S4).

**Table 3.**
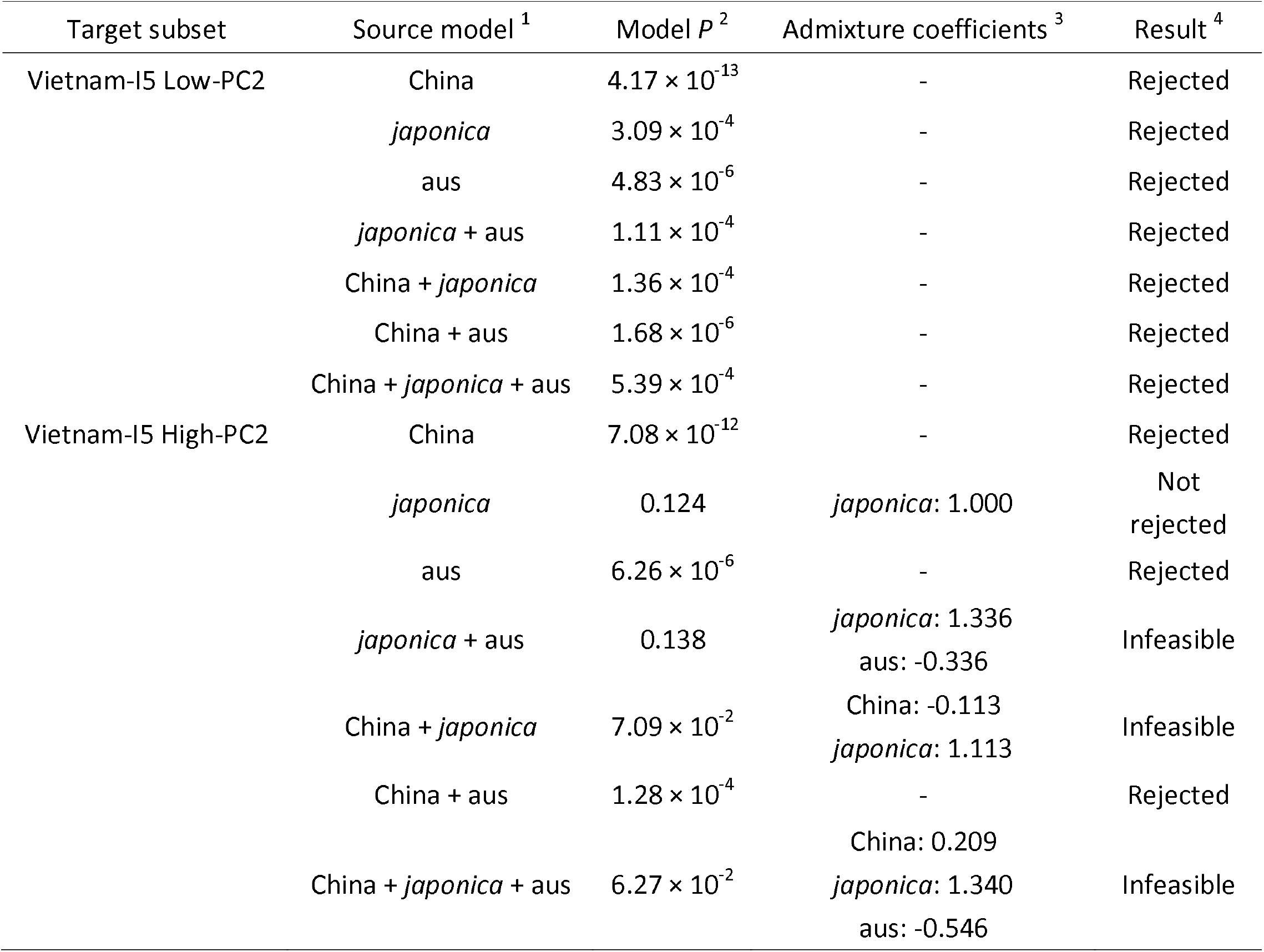
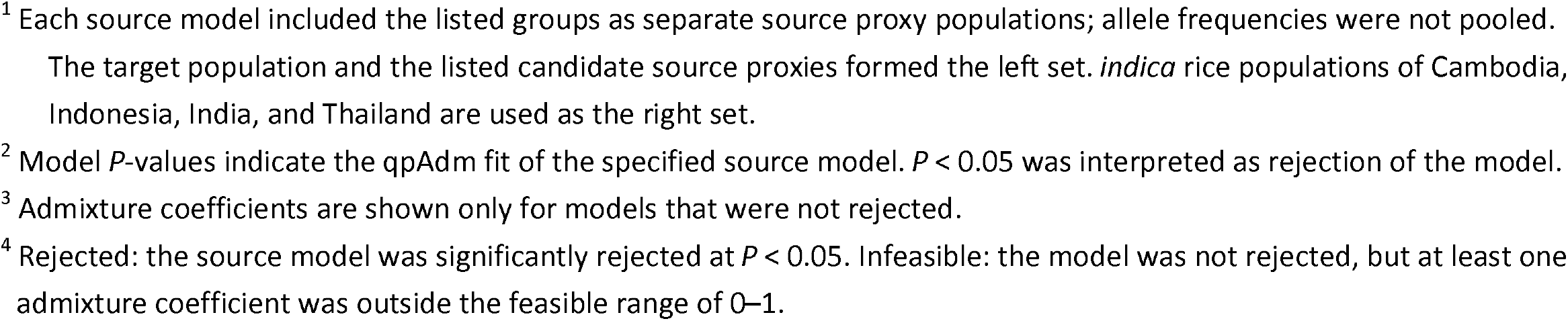
qpAdm tests of source-proxy models for the Vietnam-I5 subsets.

| Target subset | Source model <sup>1</sup> | Model $P$ <sup>2</sup> | Admixture coefficients <sup>3</sup> | Result <sup>4</sup> |
| --- | --- | --- | --- | --- |
| Vietnam-I5 Low-PC2 | China | $4.17 \times 10^{-13}$ | - | Rejected |
| | *japonica* | $3.09 \times 10^{-4}$ | - | Rejected |
| | aus | $4.83 \times 10^{-6}$ | - | Rejected |
| | *japonica* + aus | $1.11 \times 10^{-4}$ | - | Rejected |
| | China + *japonica* | $1.36 \times 10^{-4}$ | - | Rejected |
| | China + aus | $1.68 \times 10^{-6}$ | - | Rejected |
| | China + *japonica* + aus | $5.39 \times 10^{-4}$ | - | Rejected |
| Vietnam-I5 High-PC2 | China | $7.08 \times 10^{-12}$ | - | Rejected |
|  | *japonica* | 0.124 | *japonica*: 1.000 | Not rejected |
| | aus | $6.26 \times 10^{-6}$ | - | Rejected |
|  | *japonica* + aus | 0.138 | *japonica*: 1.336<br>aus: -0.336 | Infeasible |
| | China + *japonica* | $7.09 \times 10^{-2}$ | China: -0.113<br>*japonica*: 1.113 | Infeasible |
| | China + aus | $1.28 \times 10^{-4}$ | - | Rejected |
| | China + *japonica* + aus | $6.27 \times 10^{-2}$ | China: 0.209<br>*japonica*: 1.340<br>aus: -0.546 | Infeasible |

- <sup>1</sup> Each source model included the listed groups as separate source proxy populations; allele frequencies were not pooled. The target population and the listed candidate source proxies formed the left set. *indica* rice populations of Cambodia, Indonesia, India, and Thailand are used as the right set. - <sup>2</sup> Model *P*-values indicate the qpAdm fit of the specified source model. $P < 0.05$ was interpreted as rejection of the model. - <sup>3</sup> Admixture coefficients are shown only for models that were not rejected. - <sup>4</sup> Rejected: the source model was significantly rejected at $P < 0.05$ . Infeasible: the model was not rejected, but at least one admixture coefficient was outside the feasible range of 0–1.

## Discussion

Using historical landraces collected in the 1950s, this study re-examined the regional formation of *indica* lineages across East and Southeast Asia before the widespread introduction of modern cultivars. By integrating SNPs derived mainly from the intronic regions of the 66 Hamada collection accessions with a comprehensive *indica* panel compiled from previous studies (Gutaker *et al*., 2020; Higgins *et al*., 2021; Tanaka *et al*., 2020a; 2020b), we recovered the major regional clusters previously described from existing landraces while incorporating the Hamada collection into the framework (Fig. 3). These results indicate that the broad regional structure of *indica* recognized in previous studies was already in place before modern cultivar replacement and therefore reflects older historical differentiation rather than recent breeding alone.

A key outcome of this study was that north–south differentiation within Vietnam was evident in these historical accessions. Most accessions from South Vietnam, Cambodia, and Thailand were positioned around the KHM (mainly Cambodian) and THA (mainly Thai) clusters of Gutaker *et al*. (2020) but did not form sharply discrete geographic groups. Instead, they overlapped extensively with one another (Fig. 3A and S6; Table 2). The low pairwise differentiation among Cambodian, South Vietnamese, and Thai accessions further supports the view that these landraces were maintained within a connected mainland Southeast Asian agricultural network, rather than as sharply partitioned national groups. This connectivity may have been facilitated by shared rainfed lowland rice ecosystems across Mekong-associated landscapes and by historical links within the Khmer/Angkorian sphere.

In contrast, the North Vietnamese accessions formed a distinct group (Fig. 2), which is consistent with earlier observations of the distinctiveness of northern Vietnamese rice landraces (Hiraoka *et al*., 2009). Within this group, we detected two differentiated components corresponding to fifth- and tenth-month rice. This pattern is consistent with the possibility that the seasonal differentiation of northern Vietnamese rice is accompanied by an underlying genetic differentiation. In northern Vietnam, two rice-growing seasons have traditionally been recognized: tenth-month rice, sown around the fifth month of the Annamese calendar and harvested around the tenth month, and fifth-month rice, sown around the tenth month and harvested around the fifth month (Hamada, 1965). Tenth-month rice accounted for approximately 70% of the paddy rice area in northern Vietnam, whereas fifth-month rice accounted for the remaining 30% (Hamada, 1965).

The fifth-month rice accessions clustered with the Vietnam-I5 cluster, whereas the tenth-month rice accessions were not assigned to any major cluster by PAM-silhouette clustering, although both PCA and outgroup *f*_3_ indicated high affinity to the China-related group (Fig. 3). Together, these results suggest that the North Vietnamese *indica* were not represented by a single local lineage, but instead reflected multiple genetic backgrounds shaped in relation to southern China. However, these accessions were not completely isolated from the southern Vietnamese pool (Supplementary Fig. S4), supporting the interpretation that northern Vietnam as a contact zone rather than a fully discrete genetic region.

The recovery of Vietnam-I5 in historical material is one of the main findings of this study. Higgins *et al*. (2021) identified I5 as a highly distinctive outlying *indica* group among Vietnamese landraces, and our analyses recovered this lineage as the Vietnam-I5 cluster (Fig. 3). The inclusion of Hamada collection fifth-month rice accessions in Vietnam-I5 indicates that this lineage was already present before the onset of modern breeding. Its formation history appears more complex than that of a simple derivative of the Chinese cluster. Admixture graph and TreeMix analyses were both consistent with a model in which Vietnam-I5 is based on a China-related lineage but also includes an additional component (Fig. 3C; Supplementary Fig. S7). Although Vietnam-I5 showed strong affinity to the China cluster, it was separated from China primarily along PC2, and High-PC2 accessions became progressively more distant not only from China, but also from the other *indica* clusters (Figs 3, 4D,E). qpAdm showed that all tested source-proxy models were rejected for the Low-PC2 subset. These included the one-source models (China only, *japonica* only, and aus only), two-source models (*japonica* + aus, China + *japonica* and China + aus), and the three-source model (China + *japonica* + aus) (Table 3; Supplementary Table S4). For the High-PC2 subset, however, the *japonica*-only model was not rejected when only *indica* populations were used as right-set references, but this model was strongly rejected when *O. barthii* was added to the right-set (Table 3; Supplementary Table S4). Therefore, the High-PC2 pattern is consistent with elevated *japonica*-related affinity, but does not support a simple *japonica*-only origin or allow us to infer a specific *japonica* introgression event.

Together, these results indicate that Vietnam-I5 cannot be adequately explained by the sampled China-, *japonica*-, and aus-related source proxies across the tested right-set configurations. In combination with the admixture graph results (Fig. 3), these findings suggest that Vietnam-I5 may have a China-associated *indica* background, with the High-PC2 subset showing *japonica*-related affinity, while its ancestry relationships are not fully represented by the reference groups analyzed here.

This pattern is consistent with a broader historical interpretation in which northern Vietnam functioned as a zone of long-term interaction between lineages associated with southern China and those of mainland Southeast Asia. Archaeobotanical studies suggest that the spread of agriculture from southern China to mainland Southeast Asia proceeded through multiple pathways, with northern Vietnam representing an important zone of contact (Chi and Hung, 2010). Northern Vietnam also experienced prolonged political and economic connections with the Chinese world (Britannica, “Vietnam under Chinese rule”, https://www.britannica.com/place/Vietnam/Vietnam-under-Chinese-rule; accessed on April 17, 2026). Against this background, the close relationship between North Vietnamese groups and China-related lineages is not surprising. In parallel, the strong assotiation of South Vietnamese accessions to Cambodia/Thailand-related lineages indicates that southern Vietnam was part of the *indica* diversity maintained within the Mekong-linked mainland Southeast Asian agricultural sphere. Vietnam should therefore be viewed not as a region occupied by a single *indica* population, but rather as a zone in which multiple lineages co-exist and are repeatedly reshaped through interactions with both the Chinese world in the north and the Mekong world in the south. This interpretation is consistent with the strong internal structure of Vietnamese landraces reported by Higgins *et al*. (2021). It adds historical depth to the broader eastward expansion framework proposed by Gutaker *et al*. (2020) (Fig. 5).

**Fig. 5.**
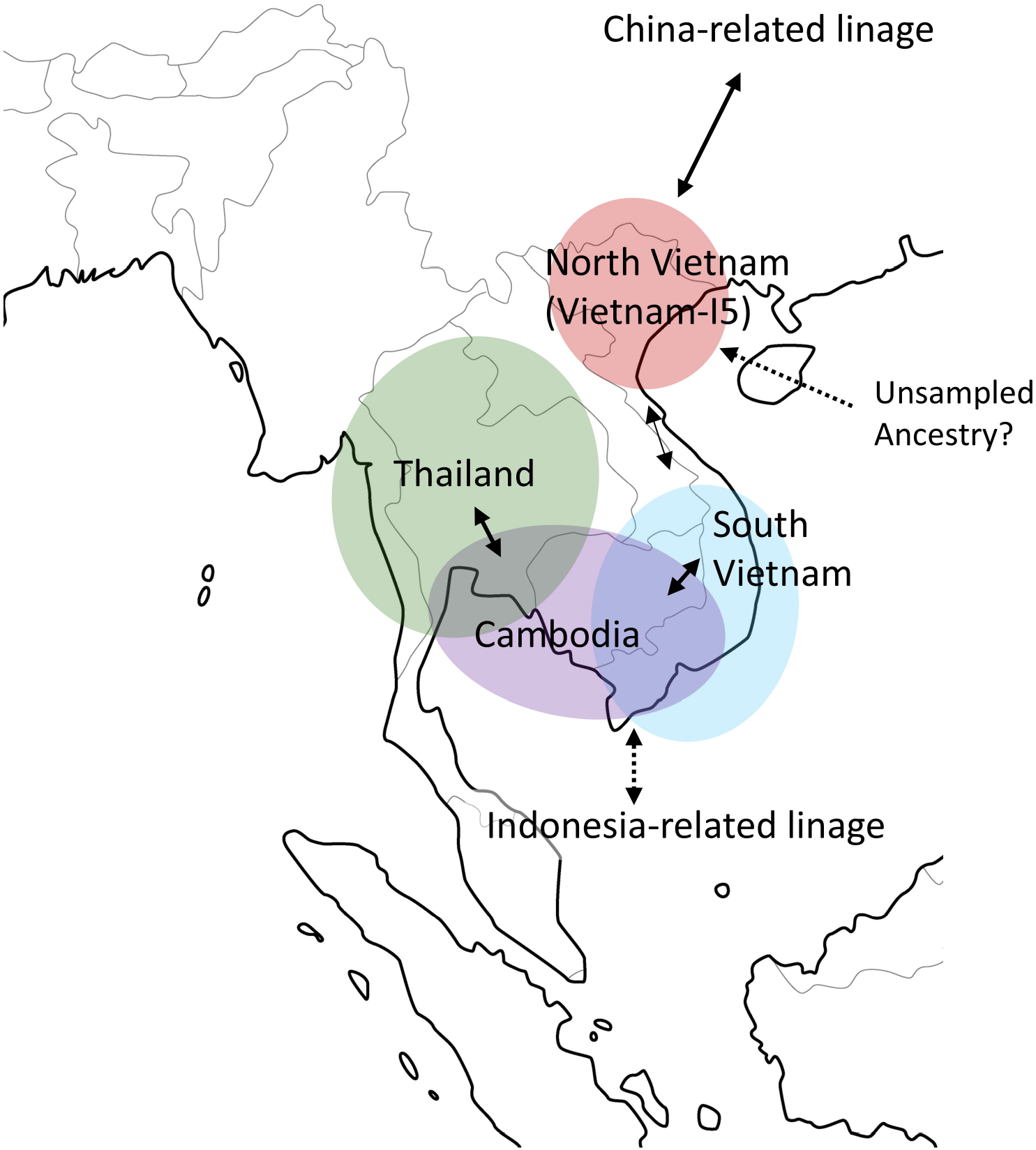
Schematic model of the historical formation of *indica* lineages in mainland Southeast Asia. Ellipses indicate approximate genetic affinities and overlap among regional lineage groups, rather than precise geographic distributions. Solid arrows indicate relationships supported by the present analyses, whereas dotted arrows indicate inferred but unresolved relationships. The arrows represent inferred contributions to affinity or ancestry and do not imply direction-specific migration routes.

We note that our admixture graph (Fig. 3C) was used as a simplified model of allele-sharing relationships among selected populations. It was not intended to reconstruct the deep dispersal history of rice. Therefore, we did not interpret the relative placement of the Chinese and Indian clusters in the graph as evidence of the timing or direction of the divergence between these lineages. Instead, we restricted our interpretation to the local relationships involving Vietnam-I5, the Chinese cluster, and Southeast Asian groups.

The main contribution of this study is the use of historical DNA to show that the regional population structure of *indica* observed in previous studies was established before the introduction of modern breeding. However, this study had certain limitations. The sample size was modest, and targeted resequencing did not permit a full evaluation of fine-scale introgression, haplotype-level formation history, or structural variation. The Hamada collection contains more than 1,000 additional accessions, some of which are reported to have been transferred to the National Institute of Genetics, Japan (Lim *et al*., 2024, 2022). Expanded whole-genome sequencing of this material should provide a much higher resolution of the regional differentiation within Vietnam, the formation of China-related lineages, and the genomic contributions of other cultivated/wild ancestry sources. Such work would not only deepen our understanding of lineage formation in pre-modern Asian rice cultivation, but also help to reposition the diversity preserved in historical landraces as a valuable resource for future rice improvement.

## Supplementary data

**Fig. S1:** MapDamage2 summary plots for the historical DNA library of ViS-956.

**Fig. S2:** PAM-silhouette clustering results for Panel 1 at Kd values from 2 to 10.

**Fig. S3**: ADMIXTURE analysis for Hamada collection populations (Panel 2).

**Fig. S4:** TreeMix models for Hamada collection populations (Panel 2).

**Fig. S5:** PAM-silhouette clustering results for Panel 1 at Kd values from 2 to 10.

**Fig. S6:** Principal component analysis (PCA) plots colored according to population assignments in each reference panel.

**Fig. S7:** TreeMix models for *indica*-wide discrete subpopulations (Panel 2).

**Table S1:** Materials used and sequencing read statistics of the Hamada collection.

**Table S2:** Whole genome sequence data downloaded and merged with that of the Hamada collection.

**Table S3:** Percentage of genomic locations of single-nucleotide polymorphisms (SNPs) derived from targeted resequencing.

**Table S4:** qpAdm tests of source-proxy models for the Vietnam-I5 subsets including *Oryza barthii* as an outgroup right population.

## Acknowledgements

We thank Ms. Meng-Ting Hsieh for the valuable comments on this manuscript.

## Author contributions

KN and TI planned and designed this study. KN, LS, OC, and RI carried out the targeted resequencing experiments. KN, YH, HS, YS, and RMG analyzed the data. KN, RMG, and TI wrote and revised the manuscript.

## Data availability

Raw FASTQ reads for 96 Hamada collection accessions have been deposited under the accession number PRJDB40808 (DRR932751-DRR932846). The sources of all downloaded data are provided in the Supplementary Information.

