## Supplementary figures S1-S7 for "Historical rice seeds from the 1950s reveal pre-modern genetic structure in *indica* landraces of mainland Southeast Asia"

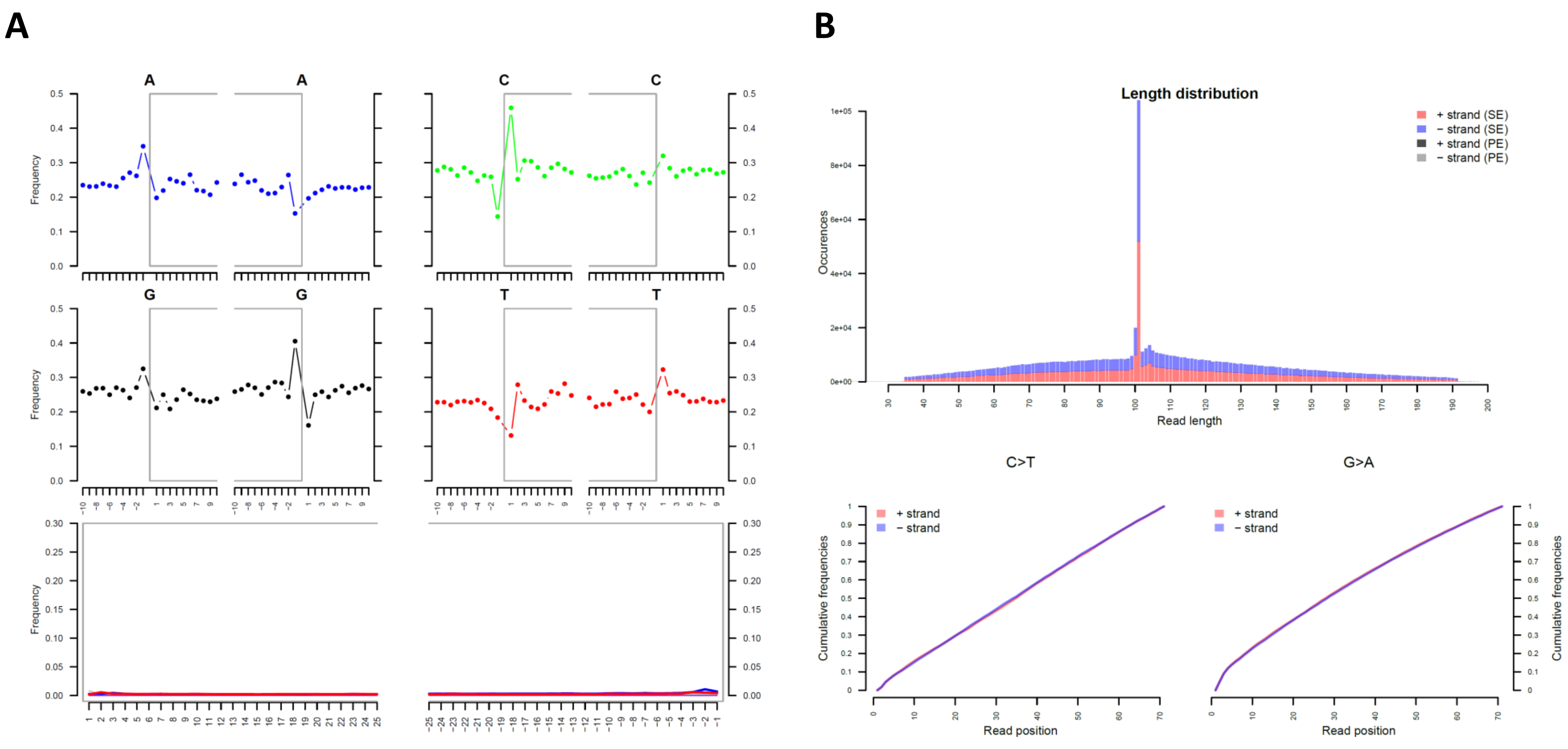


**Supplementary Fig. S1.** MapDamage2 summary plots for the historical DNA library of ViS-956.
**A** Base composition and nucleotide misincorporation patterns across read positions generated by MapDamage2. Frequencies of A, C, G, and T are plotted along read positions, with terminal substitution patterns shown for each nucleotide. **B** Read-length distribution and cumulative misincorporation patterns generated by MapDamage2. The upper panel shows read-length distributions, and the lower panels show cumulative C-to-T and G-to-A substitution frequencies along read positions.

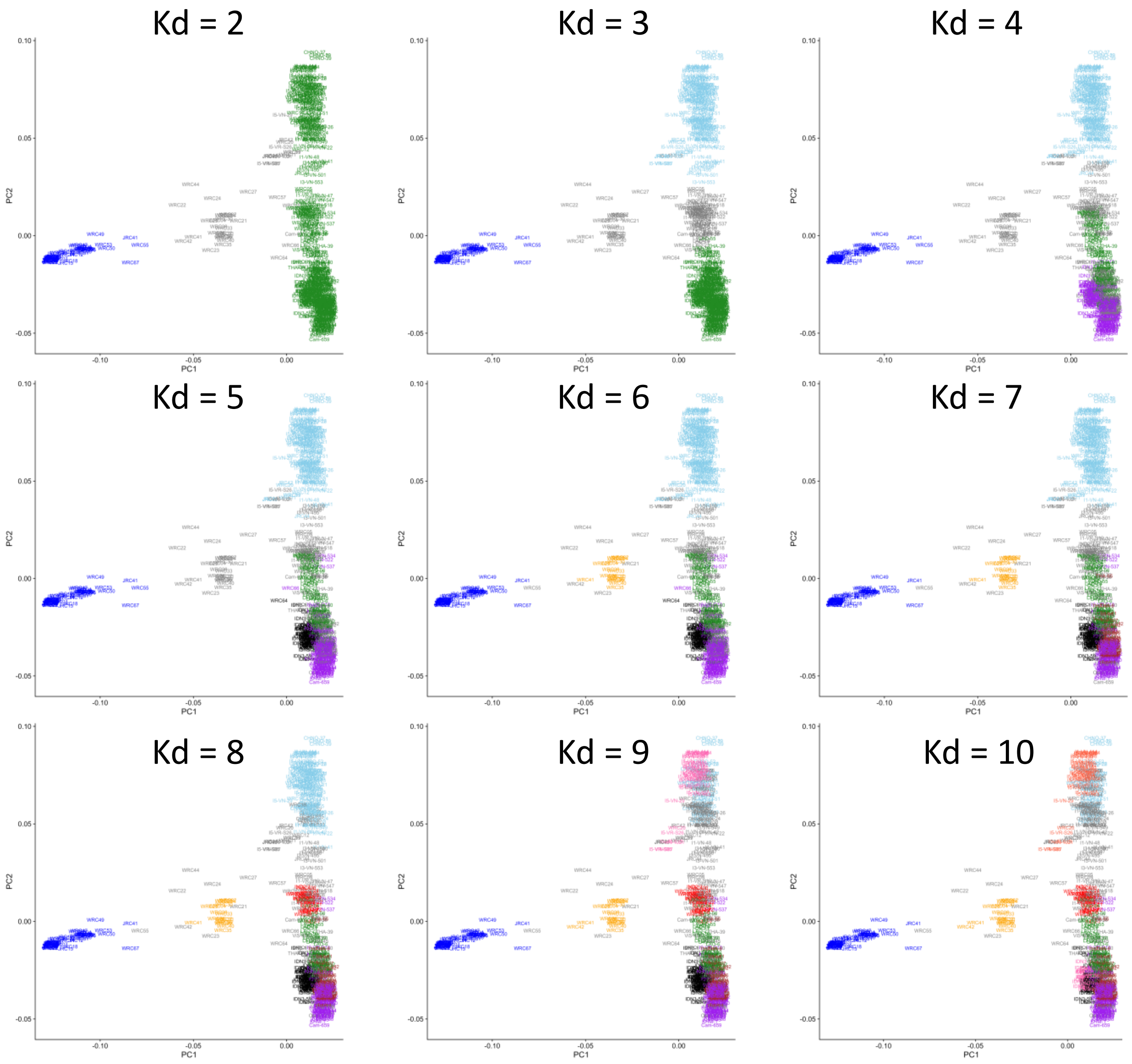


**Supplementary Fig. S2.** PAM-silhouette clustering results for Panel 1 at Kd values from 2 to 10.

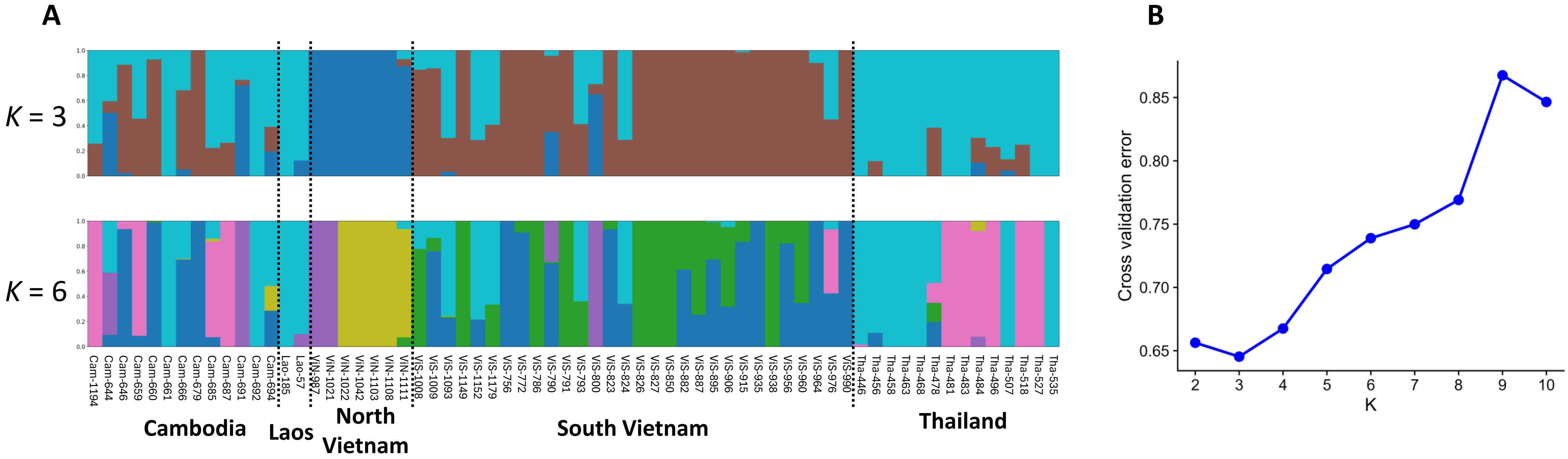


**Supplementary Fig. S3.** ADMIXTURE analysis for Hamada collection populations (Panel 2).
**A** ADMIXTURE results shown for *K* = 3 and *K* = 6. **B** Transition of cross validation error rates for the ADMIXTURE analyses.

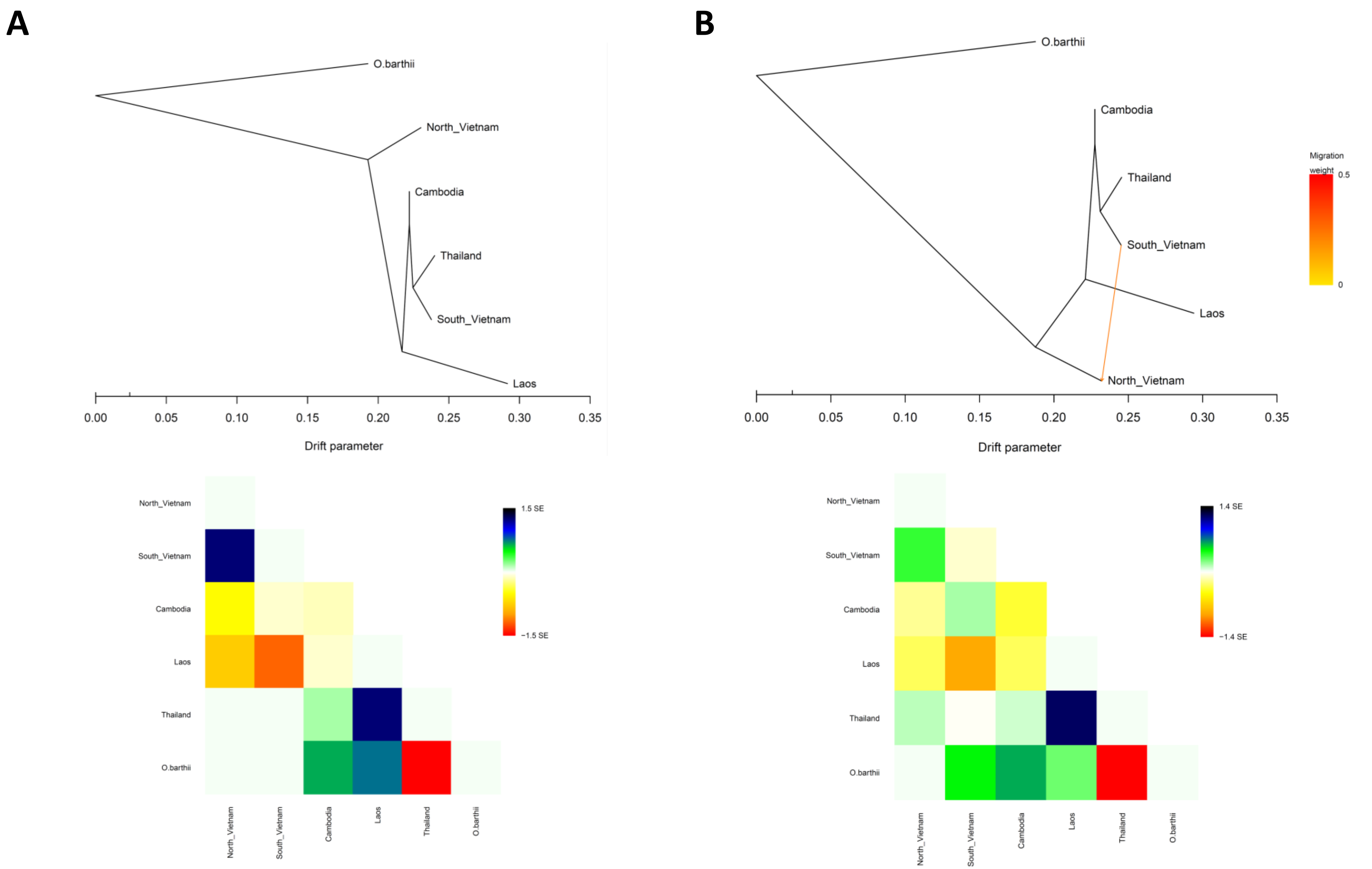


**Supplementary Fig. S4.** TreeMix models for Hamada collection populations (Panel 2).
**A** Model with no migration edge (m = 0). **B** Model with one migration edge (m = 1). The upper panels show the maximum-likelihood trees inferred by TreeMix. In **B**, the colored arrow indicates the inferred migration edge, with color intensity representing migration weight. The lower panels show the corresponding residual fit plots among the tested populations; residuals closer to zero indicate a better fit of the model to the data.

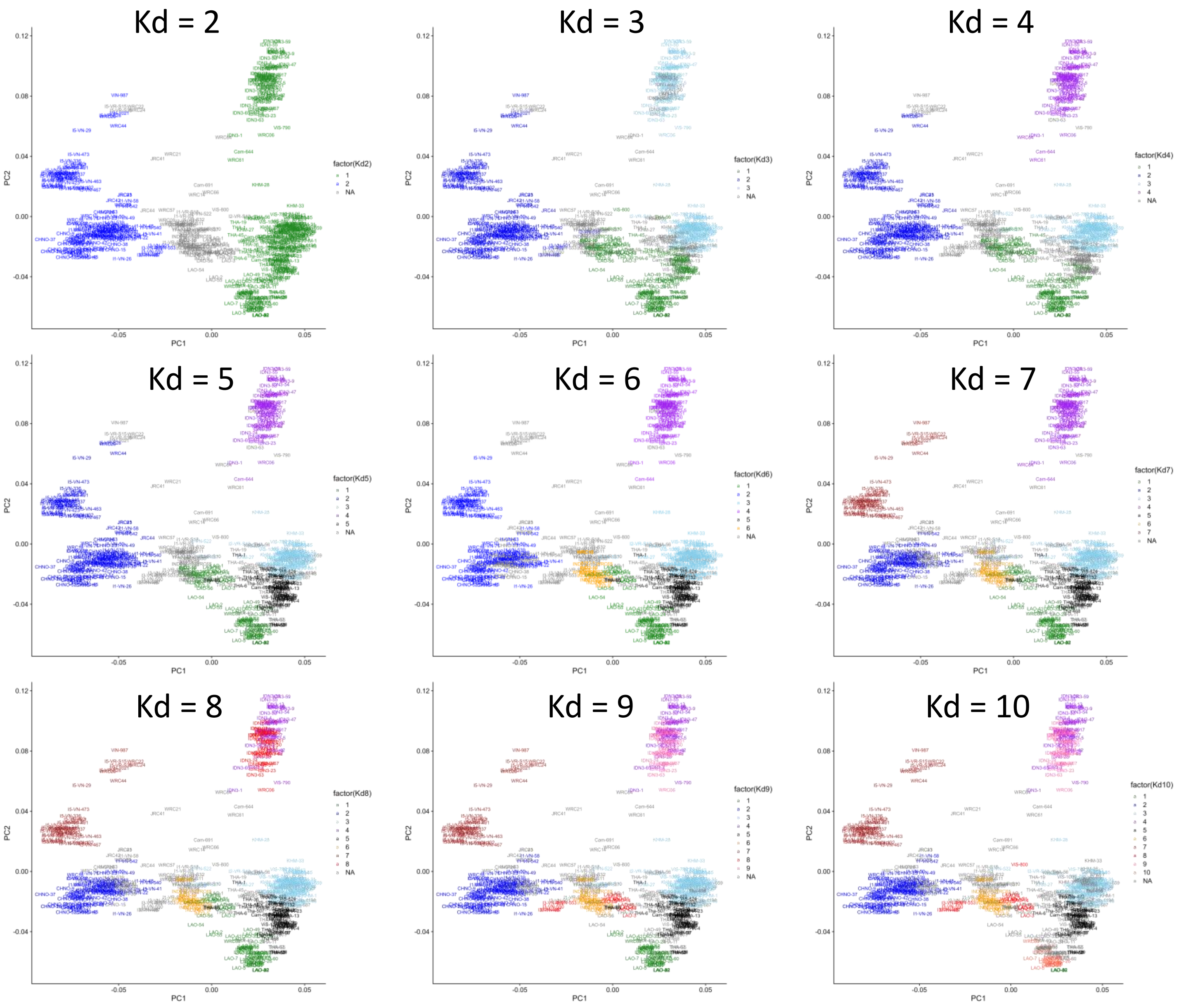


**Supplementary Fig. S5.** PAM-silhouette clustering results for Panel 3 at Kd values from 2 to 10.

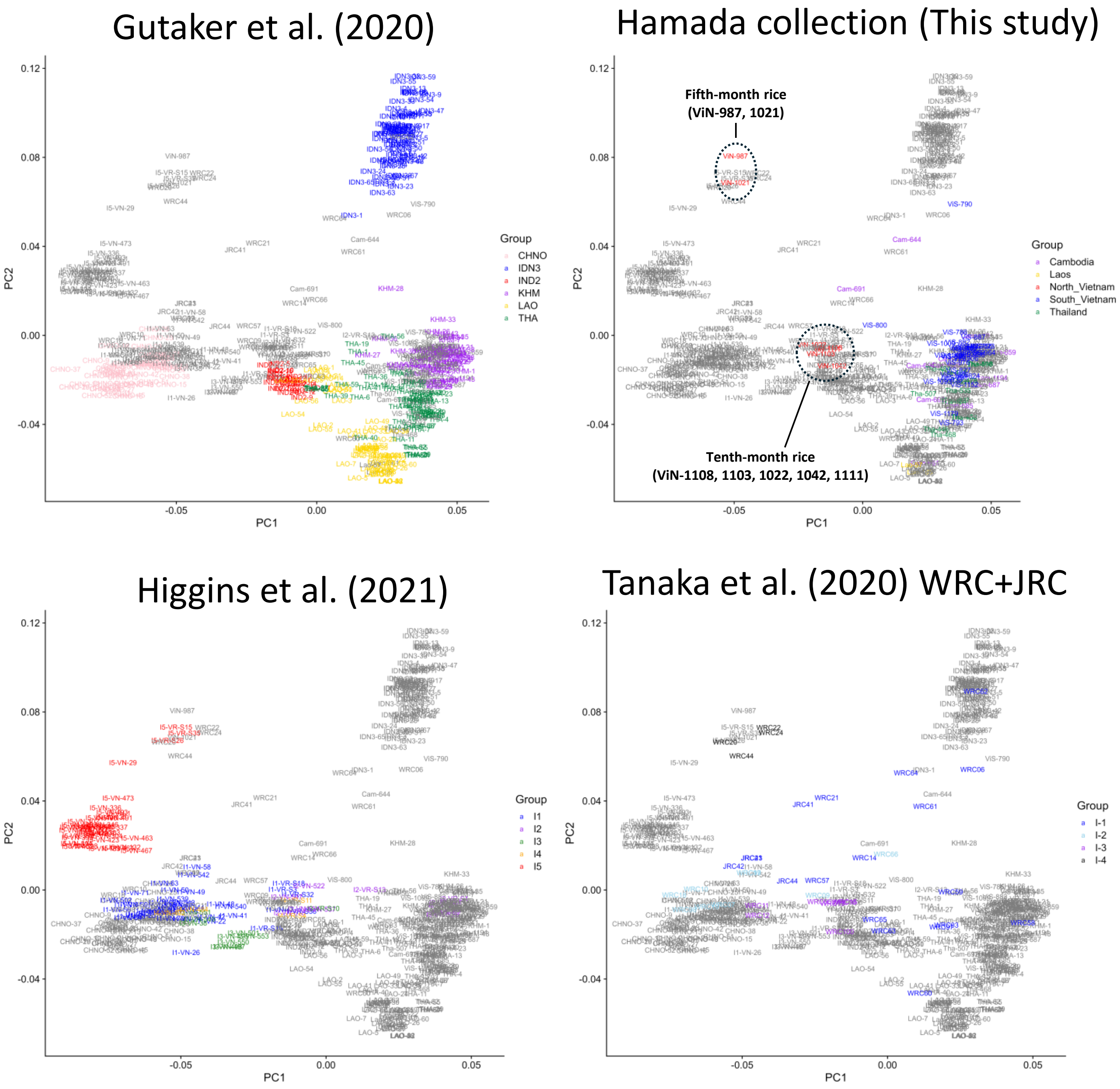


**Supplementary Fig. S6.** Principal component analysis (PCA) plots colored according to population assignments in each reference panel.

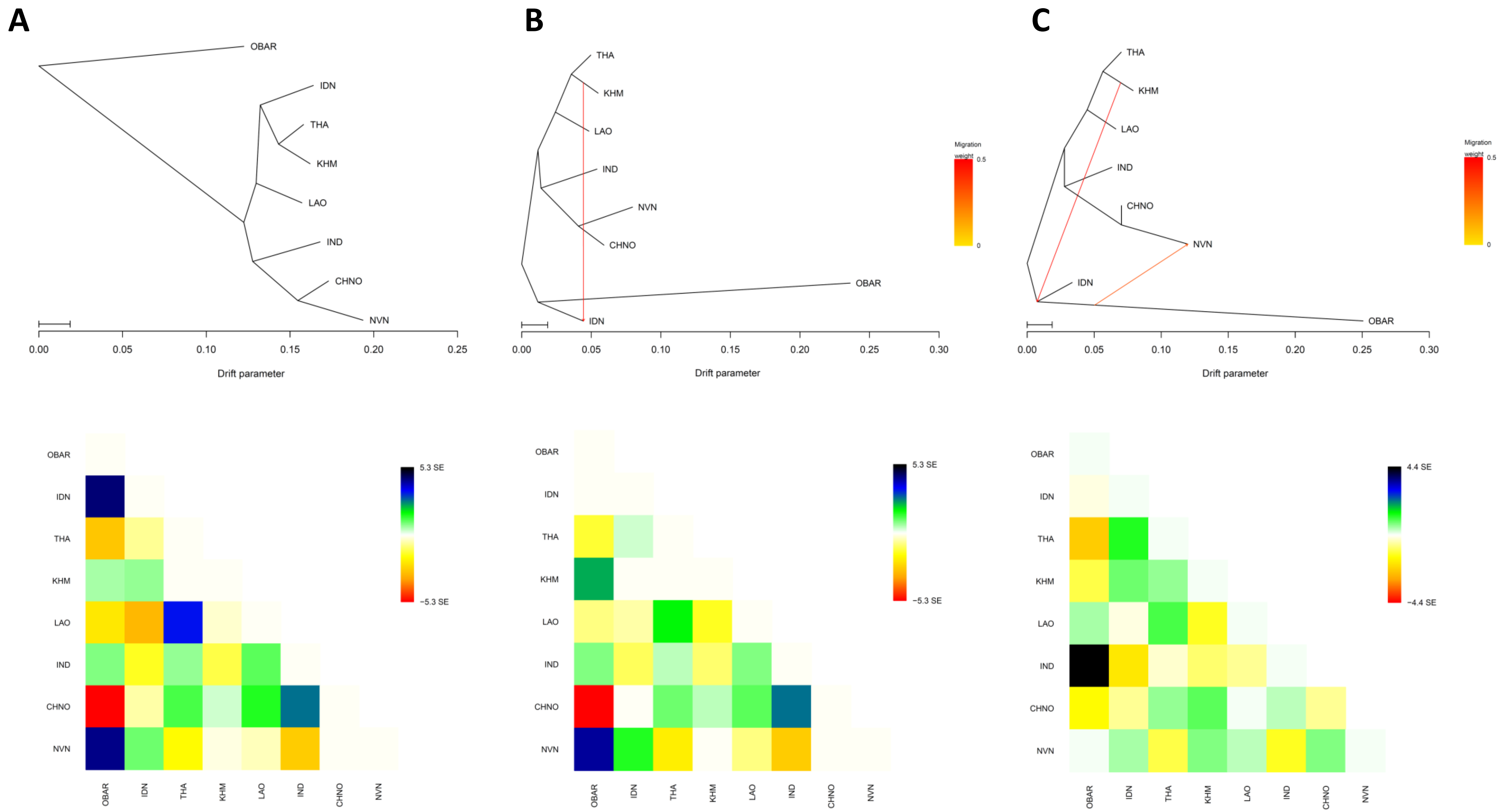


**Supplementary Fig. S7.** TreeMix models for *indica*-wide discrete subpopulations (Panel 2).
**A** Model with no migration edge (m = 0). **B** Model with one migration edge (m = 1). **C** Model with two migration edges (m = 2). The upper panels show the maximum-likelihood trees inferred by TreeMix. In **B** and **C**, the colored arrows indicates the inferred migration edge, with color intensity representing migration weight. The lower panels show the corresponding residual fit plots among the tested populations; residuals closer to zero indicate a better fit of the model to the data.
